# Differential Expression of 5’-tRNA Fragments in Circulating Preeclampsia Syncytiotrophoblast Vesicles Drives Macrophage Inflammation

**DOI:** 10.1101/2023.04.11.536371

**Authors:** William Robert Cooke, Peiyong Jiang, Lu Ji, Jinyue Bai, Gabriel Davis Jones, Y. M. Dennis Lo, Christopher Redman, Manu Vatish

**Author notes:** Corresponding author: Dr William Cooke, Nuffield Department of Women’s and Reproductive Health, University of Oxford Level 3, Women’s Centre, John Radcliffe Hospital. Oxford, *OX3 9DU*, UK.

## Abstract

**Background:** The relationship between placental pathology and the maternal syndrome of preeclampsia is incompletely characterised. Mismatch between placental nutrient supply and fetal demands induces stress in the syncytiotrophoblast, the layer of placenta in direct contact with maternal blood. Such stress alters the content and increases the release of extracellular vesicles (STB-EVs) into the maternal circulation. We have previously shown 5’-tRNA fragments (5’-tRFs) constitute the majority of small RNA in STB-EVs in healthy pregnancy. 5’-tRFs are produced in response to stress. We hypothesised STB-EV 5’-tRF release might change in preeclampsia.

**Methods:** We perfused placentas from eight women with early-onset preeclampsia and six controls, comparing small RNA expression in STB-EVs. We used membrane-affinity columns to isolate maternal plasma vesicles and investigate placental 5’-tRFs *in-vivo*. We quantified 5’-tRFs from circulating STB-EVs using a placental alkaline phosphatase immunoassay. 5’-tRFs and scrambled RNA controls were added to monocyte, macrophage and endothelial cells in culture to investigate transcriptional responses.

**Results:** 5’-tRFs constitute the majority of small RNA in STB-EVs from both preeclampsia and normal pregnancies. >900 small RNA fragments are differentially expressed in preeclampsia STB-EVs. Preeclampsia-dysregulated 5’-tRFs are detectable in maternal plasma, where we identified a placentally-derived load. 5’-tRF-Glu-CTC, the most abundant preeclampsia-upregulated 5’-tRF in perfusion STB-EVs, is also increased in preeclampsia STB-EVs from maternal plasma. 5’-tRF-Glu-CTC induced inflammation in macrophages but not monocytes. The conditioned media from 5’’-tRF-Glu-CTC-activated macrophages reduced eNOS expression in endothelial cells.

**Conclusions:** Increased release of syncytiotrophoblast-derived vesicle-bound 5’-tRF-Glu-CTC contributes to preeclampsia pathophysiology.

## Introduction

Preeclampsia is a complex placental syndrome, with multiple causes and a variable phenotype. Clinical features are characterised by maternal sterile inflammation and endothelial dysfunction^1^. A point of convergence in the disorder is stress in the syncytiotrophoblast, the interface between maternal and fetal circulations^2^. The link between syncytiotrophoblast stress and maternal symptoms, likely through blood-borne factors, is incompletely characterised^3^. Angiogenic proteins (soluble fms-like tyrosine kinase [sFlt-1] and placental growth factor [PlGF]) are important syncytiotrophoblast stress signals which contribute to the preeclamptic syndrome^4^. Excess sFlt-1 sensitises endothelial cells to pro-inflammatory cytokines^5^. These molecules have been successfully applied to the clinical diagnosis of preeclampsia^6^. Altered extracellular vesicle [EV] release from the syncytiotrophoblast, whilst also a key contributor in the pathophysiology of preeclampsia, remains underexplored^7^.

The healthy syncytiotrophoblast releases EVs [STB-EVs] directly into the maternal circulation^7^. These lipid bilayer-bound particles are decorated with surface proteins and shuttle their contents to distant cells. Cellular stress increases EV release; this is reflected in preeclampsia where circulating STB-EVs are more abundant^8^. STB-EV cargoes also change in preeclampsia; for example nitric oxide synthase expression is reduced and neprilysin (a metalloprotease causing hypertension) is increased^9,10^.

We recently reported 5’-transfer RNA fragments [5’-tRFs] as the predominant small RNA species in healthy STB-EVs^11^. 5’-tRFs form when mature tRNA molecules are cleaved by a number of stress-induced ribonucleases including angiogenin^12^. They can be exported as EV cargo^13^. 5’-tRF expression profiles can be complex; at least 417 tRNA genes can be cleaved at multiple loci^14^. 5’-tRFs are also multifaceted signalling molecules, regulating transcription, translation and epigenetic inheritance^15^. 5’-tRFs have mostly been investigated in cancer biology and immunology where they are described as intracellular, autocrine and paracrine signals^13,16,17^.

5’-tRFs are stress signals; syncytiotrophoblast stress is a key feature of preeclampsia. We hypothesised that syncytiotrophoblast 5’-tRF release may change in preeclampsia. We used placental perfusion as a source of STB-EVs to show that 5’-tRF expression in preeclampsia differed from healthy pregnancy. Our *in-vivo* work demonstrated a placentally-derived load of circulating 5’-tRFs. 5’-tRF-Glu-CTC (the most abundant preeclampsia-upregulated 5’-tRF) was increased in preeclampsia plasma STB-EVs. Cell culture studies found 5’-tRF-Glu-CTC triggered sterile inflammation in macrophages. Together these findings suggest 5’-tRFs may link syncytiotrophoblast stress with maternal inflammation in preeclampsia.

## Methods

### 1. Sample collection and storage

This project was approved by the Central Oxfordshire Research Ethics Committee (07/H0607/74 and 07/H0606/148). All participants provided informed written consent. Preeclampsia was defined using the ISSHP classification^18^. Placentas were obtained at the time of caesarean section and perfused within 10 minutes of delivery. Uterine vein samples were taken during caesarean section, just prior to uterine incision, ipsilateral to the placental site. Peripheral blood samples were taken from the antecubital fossa. 21-gauge needles and 4.5 ml sodium citrate vacutainers were used for venepuncture (BD Diagnostics, UK). Non-pregnant samples were from female volunteers of reproductive age. Plasma was obtained by centrifugation at 1500 *g* for 15 minutes. Samples were processed within 30 minutes of collection, aliquoted and stored at −80°C.

### 2. Placental perfusion

Placentas from eight women with early-onset preeclampsia and six normotensive pregnancies were perfused using a well-established dual-lobe perfusion technique^19^. Maternal perfusate was centrifuged at 10 000 *g* for 30 minutes to isolate medium-large extracellular vesicles [MLEVs]. The supernatant was centrifuged at 150 000 *g* for 2 hours to isolate small extracellular vesicles [SEVs]. Biopsies of placental tissue were taken from the maternal surface of a non-perfused lobe. EVs were characterised using Nanoparticle Tracking Analysis, transmission electron microscopy and Western blotting as described previously^19^.

### 3. RNA sequencing

RNA was isolated from MLEVs, SEVs and placental tissue using Total RNA Purification Plus Kit (Norgen Biotek Corporation, Canada). After confirming RNA quantity and integrity using Bioanalyser (Agilent Technologies, Germany), the same amount of input RNA was loaded for library preparation using the NEBNext Multiplex Small RNA library preparation kit (New England Biolabs, USA). Libraries were size-selected for fragments 15-50 bp by gel electrophoresis; fragment size and concentration was confirmed using High Sensitivity D1000 ScreenTape (Agilent, UK). Single-end sequencing by synthesis was undertaken using an Illumina HiSeq 2500 machine (Illumina, USA). One preeclampsia sample was removed from the SEV/placenta groups due to a technical issue.

Sequence reads were analysed using sRNAnalyzer^20^. Briefly, sequencing adaptors and low-quality reads were removed using Cutadapt^21^. All identical reads in sequence reads were collapsed, thus generating a set of unique reads (referred to as fragment ID in this study). The number of sequence reads attributed to a fragment ID were defined as the raw expression level of such a fragment ID. The fragment IDs were mapped to the human small RNA databases allowing 2 mismatches. The human small RNA databases comprised miRNA, piRNA, snoRNA, rRNA, and tRNA^20,22,23^. The normalised expression level for each fragment ID within a class of RNA was used for downstream differential expression analyses. The normalised expression level for a fragment ID in a RNA class was calculated by dividing the raw expression level by a per million scaling factor of total reads of that RNA class, expressed as reads per million (RPM). Compared to healthy placentas, differentially expressed fragment IDs in the MLEVs of preeclampsia placentas were determined using the following criteria: i) At least one MLEV sample had expression level of greater than 100 RPM, ii) P-value was required to be less than 0.05 based on Mann Whitney test after Benjamini-Hochberg correction. The in-house bioinformatics pipeline was written in Perl and R languages for counting sequence reads, tag annotations, and differential expression analysis. Sequence read archive data for blood cell datasets were downloaded from NCBI, with identifiers shown in Supplementary Table 3.

### 4. Plasma EV RNA isolation

Plasma aliquots were thawed at 37°C and centrifuged at 3000 *g* for 5 minutes to remove cryoprecipitates. For membrane-based affinity isolation of total vesicular RNA, 500 µL plasma was loaded onto exoRNeasy™ Midi columns (Qiagen, Germany). For magnetic-bead isolation of STB-EV RNA, 500 µL plasma was centrifuged at 10 000 *g* for 30 minutes and the total EV pellet washed once before resuspension with biotin-saturated MojoSort streptavidin magnetic nanobeads (Biolegend, USA) to deplete non-specific binding. The supernatant was resuspended with nanobeads pre-bound with biotinylated in-house placental alkaline phosphatase (PLAP) antibody, known as NDOG2^24^. Bead-STB-EV complexes were washed four times before EV RNA isolation using Trizol LS (Invitrogen, USA). RNA was stored in aliquots at −80°C.

### 5. RT-qPCR detection

Custom Taqman™ stem-loop assays were designed for specific small RNA target sequences identified from RNA sequencing analysis (Applied Biosystems, USA). Assay linearity and specificity were verified. Perfusion samples were normalised to *TBP* (confirmed empirically to be a suitable reference). Plasma samples were normalised to *C. Elegans* miR-39 spike-in. qPCR assays are documented in Supplementary Table 4. QuantStudio™ qPCR instruments (Applied Biosystems, USA) automatically determined quantification cycles (Cq) using standard settings; Cq >35 was considered undetectable. Relative expression was determined by following the PFAFFL approach, normalising to median expression in the control group.

### 6. Cell culture

THP-1 cells (ATCC, USA) were seeded onto 24-well Nunc plates at 50 000 cells per well in RPMI-1640 medium supplemented with 10% Fetal Calf Serum. Cells were grown with 6.2 ng/mL phorbol 12-myristate-13-acetate (PMA) for 24 hours to differentiate into macrophages. Transfection experiments were performed using RNA oligonucleotides (IDT, USA) with a 5’-P modification; the sequence for tRF-A is shown in Supplementary Table 2. The scramble RNA control was the most abundant STB-EV tRF without 5’-P modification and U nucleotides replaced with A (sequence 5’-GCAAAGGAGGAACAGAGGAAGAAAACACGCCA-3’). 9.2uM RNA was packaged into lipid vesicles using N-[1-(2,3-Dioleoyloxy)propyl]-N,N,N-trimethylammonium methylsulfate [DOTAP], following manufacturer’s instructions (Roche, Switzerland) and in line with a prior publication^17^. Human Umbilical Vein Endothelial Cells were purchased and seeded onto 24-well Nunc plates at 25 000 cells per well in Endothelial Cell Growth Medium 2 (PromoCell, Germany). Macrophage supernatant experiments were conducted by preparing 2X Growth Medium and mixing 50:50 with THP-1 supernatant. Cellular RNA was isolated using RNeasy™ Plus Mini Kit (Qiagen, Germany). Target mRNA expression was quantified and normalised to *GAPDH.* Pre-amplification was used to detect IL-12B using TaqMan™ master mix #4391128 with 10 cycles.

### 7. Data presentation

All data within this manuscript are derived from distinct samples. Figures 3B and 4a were created using BioRender.com (Toronto, Canada). Statistical analyses and figures were generated using RStudio (RStudio, USA) and Prism9 (GraphPad, USA). Suspected outliers were excluded if over the 85^th^ centile. Unpaired two-tailed Mann Whitney tests were used throughout unless specifically stated within the figure legend.

### 8. Data availability

Data supporting findings of this study and bioinformatic pipelines are available through collaboration upon reasonable request to the corresponding author.

## Results

### The preeclamptic syncytiotrophoblast exports 5’-tRFs in EVs, mirroring healthy pregnancy

We isolated SEVs and MLEVs, alongside placental biopsies, from eight placentas with early-onset preeclampsia and six normotensive controls using dual-lobe placental perfusion. Pregnancy characteristics are shown in Supplementary Table 1. We performed single-end small RNA sequencing (size selecting <50 nucleotides). Sequence length distribution plots confirmed the majority of reads in MLEVs and SEVs from both preeclampsia and normal placentas were 30-34 nucleotides long (Figure 1A). In contrast a peak at 22 nucleotides was greater in the placental samples from both groups. The majority of fragments in MLEVs and SEVs mapped to tRNA species, rather than to the ribosomal RNA and micro-RNA species seen in placental tissues (Figure 1B). Coverage plots demonstrated that 5’-tRFs (but not 3’-tRFs) constitute almost all tRNA reads in EVs in preeclampsia as well as normal pregnancies (Figure 1C).

**Figure 1.**
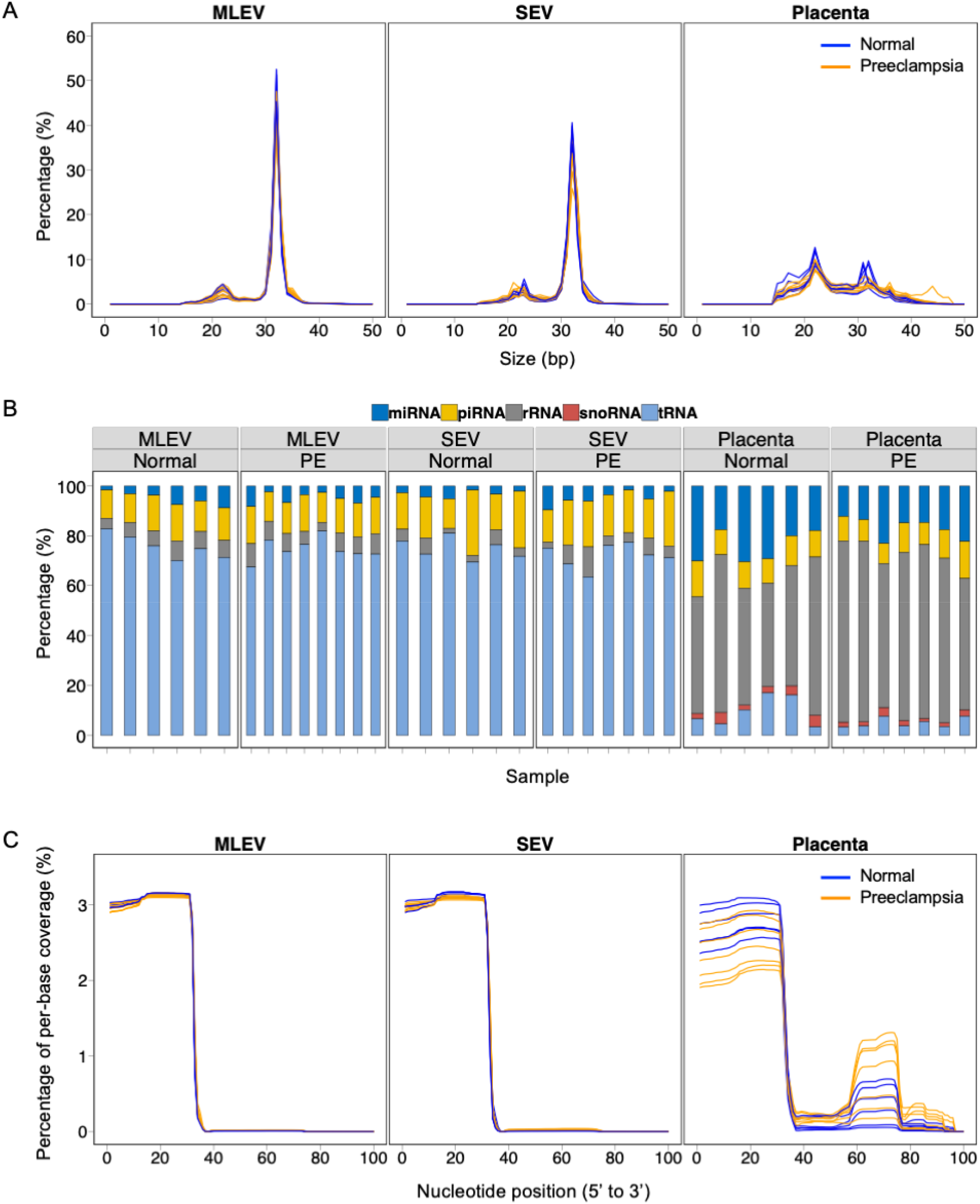
Assessment of reads <50 base pairs from medium-large extracellular vesicles (MLEV), small extracellular vesicles (SEV) and placental samples obtained from placentas in normotensive pregnancy (Normal, n=6) and early-onset preeclampsia (PE, n=8). A: Sequence length distribution plots showing size in base pairs (bp) for small RNA fragments after removal of adaptors and low-quality reads. B: Mapping of small RNA fragments. C: Coverage plots showing percentage of per-base coverage for mapped tRNA fragments.

### 5’-tRFs are differentially expressed in STB-EVs from preeclampsia compared to normotensive placentas

We used a bespoke bioinformatics pipeline to investigate differential expression of tRFs. To minimise data loss, we assigned each unique fragment an identifier, then annotated fragments after differential expression analysis (Methods). We identified 983 differentially expressed small RNA fragments in preeclampsia MLEVs compared to controls; 182 mapped to 5’-tRFs using GtRNAdb 2.0^23^. No fragments were found to be differentially expressed in preeclampsia SEVs.

The 12 most abundant differentially expressed fragments are shown in Table 1. Using the sum of preeclampsia MLEV normalised counts as a denominator, these 12 fragments account for 64% of the differentially expressed counts. 5% of these counts were accounted for by the 626 least abundant fragments. Thus, a small number of abundant fragments represented the majority of the signal in an otherwise complex dysregulated small RNA profile in preeclampsia MLEVs.

**Table 1.**
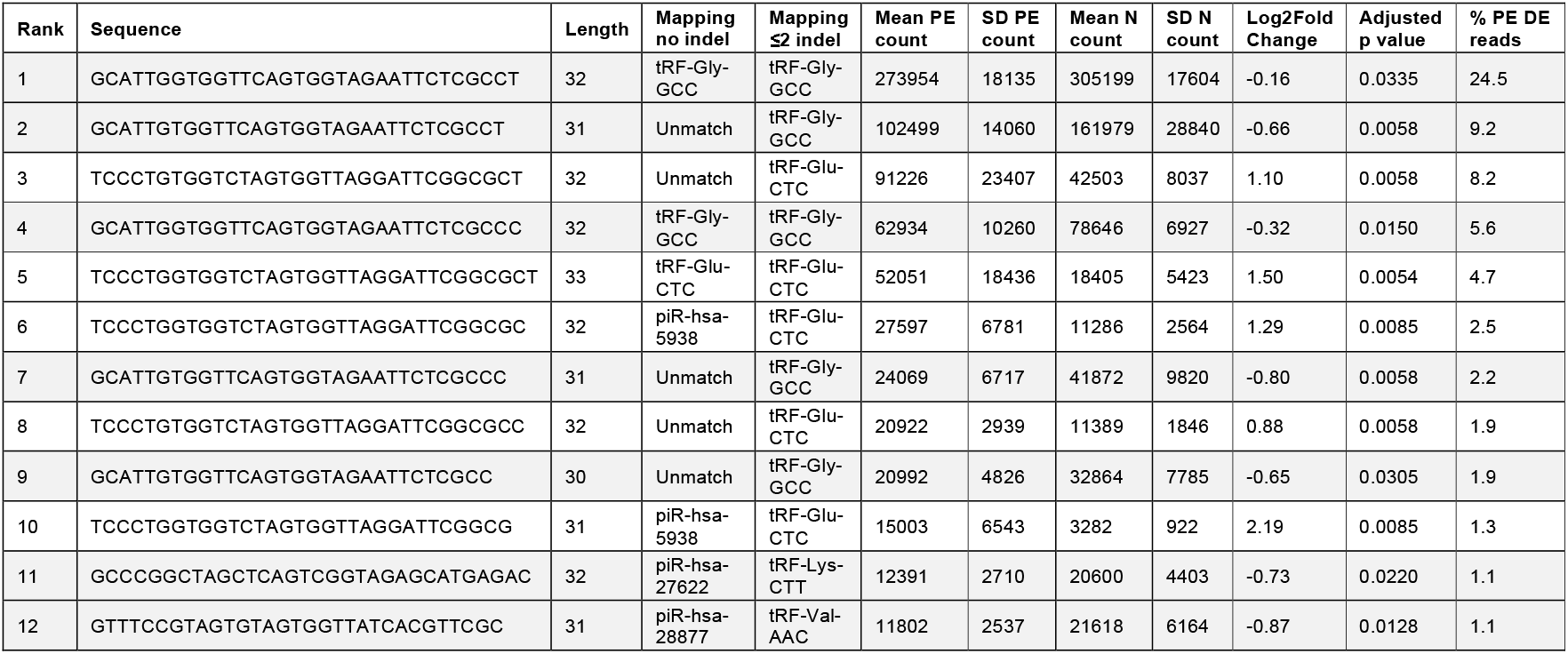
Descriptors of twelve most abundant differentially expressed small RNA fragments in early-onset preeclampsia (PE) perfusion STB-EVs compared to normotensive (N) perfusion STB-EVs, ranked by abundance. Rows showing targets downregulated in preeclampsia are shaded grey. Mapping is shown with no insertions or deletions (indel) or permitting up to two. Standard deviation (SD), differentially expressed (DE).

Comparing to human small RNA databases using conventional mapping, among the 12 most abundant differentially expressed fragments, 3 were identified as 5’-tRFs and 4 as piwi-interacting RNAs. The remaining 5 had no directly matched reference sequences. Review of fragment sequences demonstrated substantial overlap; indeed all 12 showed only one or two nucleotide insertions or deletions differentiating them from known 5’-tRFs (Table 1). By adapting the post-hoc mapping strategy to incorporate up to two insertions or deletions, it was evident that the majority of differentially expressed small RNA in preeclampsia are 5’-tRFs. A complex profile of differential expression in preeclampsia MLEVs was thus dominated by minor variants of a handful of abundant 5’-tRFs.

We sought three target 5’-tRFs which were differentially expressed in preeclampsia STB-EVs to validate our findings *ex-vivo* (in placental perfusion samples) and *in-vivo* (in plasma). 5’-tRFs are known to be expressed in other circulating EVs, most notably from immune cells. The most abundant EVs in plasma are blood-cell derived. Hence, we used publicly available blood-cell datasets to quantify possible target 5’-tRFs likely to be within contaminating EVs in plasma. Using these data we selected three tRFs for validation which were: abundant in STB-EVs (above 90^th^ centile PE expression amongst 983 differentially expressed fragments), upregulated in preeclampsia, and of low relative abundance in potentially contaminating EV source cells (Supplementary Table 2). The relative abundance of these three targets in STB-EVs using RNA sequencing is shown in Figure 2A (5’-tRF targets named A, B and C for brevity but full sequences and tRNA derivations shown in Supplementary Table 2).

**Figure 2.**
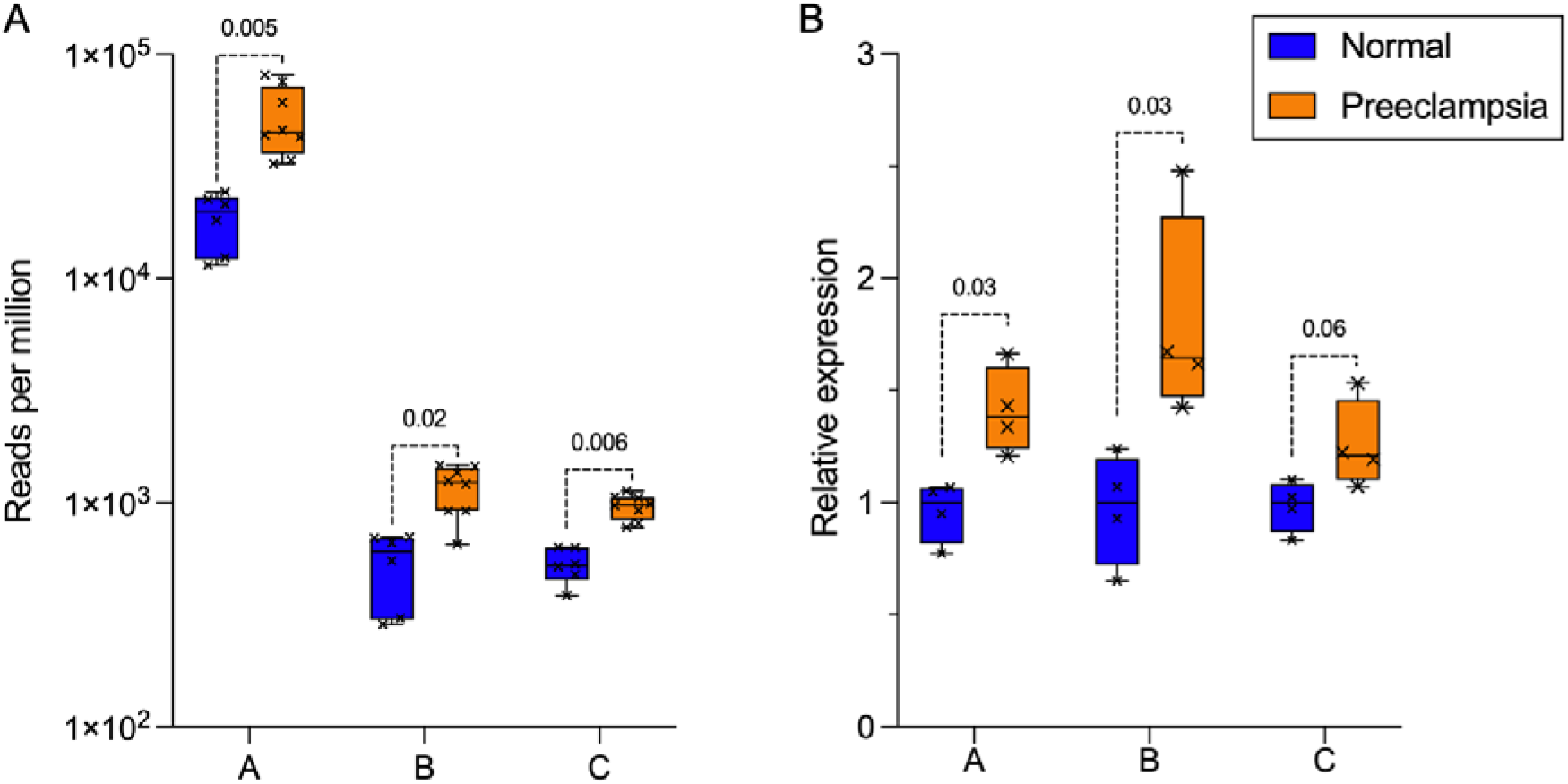
Expression of three target small RNA fragments in early-onset preeclampsia and normotensive pregnancy STB-EVs obtained by placental perfusion. A: Quantified using RNA sequencing (n=6 normotensive, n=8 preeclampsia; Mann Whitney tests after Benjamini-Hochberg correction displayed). B: Quantified using stem-loop qPCR (normalised to reference gene TBP, n=4 per group). Boxes show median/interquartile range; whiskers show max/min. Fragments labelled A, B, C for brevity; full sequences in Supplementary Table 2.

We used custom small RNA assays with stem-looped reverse transcription primers and specific minor-groove binding TaqMan™ probes to compare relative 5’-tRF expression in STB-EVs obtained by perfusion using quantitative real-time polymerase chain reaction. Results validated target 5’-tRF upregulation and effect sizes were consistent with RNA sequencing findings (median 1.4-fold upregulated in preeclampsia, Figure 2B).

### A proportion of 5’-tRFs are pregnancy-specific and placentally-derived in the maternal circulation

Total EV RNA was isolated from maternal plasma using membrane-affinity columns. EV size profiles for plasma were comparable to perfusion MLEVs (Supplementary Figure 1). Extensive characterisation of plasma membrane-affinity EVs has previously been published^25^. Target 5’-tRFs were significantly more abundant in pregnant peripheral plasma EVs than in non-pregnant matched control samples (median 3.0-fold higher in pregnancy, p<0.05 for all three) (Figure 3A). The difference between these sample groups suggested a pregnancy-specific load of 5’-tRFs in plasma EVs. The high abundance of these 5’-tRFs in non-pregnant samples confirms they were not unique to pregnancy (median Cq values in non-pregnant samples: A 24.0, B 27.9, C 25.9). We then acquired paired plasma samples simultaneously from the uterine and peripheral veins of women without preeclampsia undergoing elective caesarean section for an indication unrelated to preeclampsia (e.g. breech presentation) prior to delivery of the feto-placental unit. The uterine vein directly receives blood from the placenta, thus placentally-derived molecules are more abundant in these samples (Figure 3B)^26^. All three 5’-tRFs were more abundant in uterine vein samples (median 1.3-fold, p<0.05 for all three) (Figure 3C). These data support a placentally-derived load of 5’-tRFs in circulating plasma EVs.

**Figure 3.**
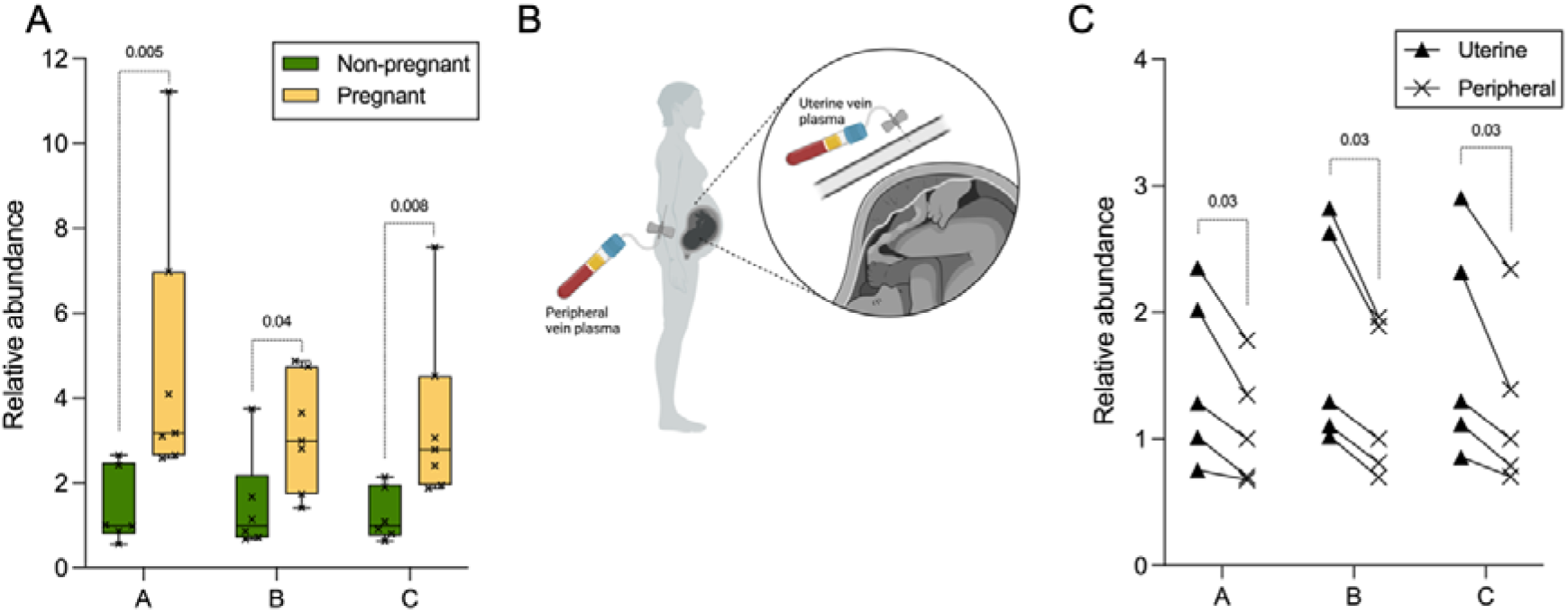
Detection of three target small RNA fragments in plasma using qPCR A: Expression in total EVs isolated from peripheral venous plasma from seven healthy third trimester pregnancies and six female volunteers of reproductive age (boxes show median/interquartile range; whiskers show max/min). B: Diagram demonstrating sampling rationale for uterine and peripheral venous blood C: Expression in total EVs isolated from uterine and peripheral venous plasma taken simultaneously from five healthy term pregnancies (paired one-tailed Wilcoxon tests displayed). Fragments labelled A, B, C for brevity; full sequences in Supplementary Table 2.

### STB-EV 5’-tRF-Glu-CTC, the most abundant preeclampsia-upregulated 5’-tRF, is increased in preeclampsia maternal plasma

A technique was optimised (Figure 4A) to isolate STB-EV RNA from maternal plasma, targeting the syncytiotrophoblast marker protein placental alkaline phosphatase (PLAP). Streptavidin nanobeads (130 nm diameter) were incubated with a highly specific biotinylated anti-PLAP antibody (NDOG2, in-house). PLAP+ plasma MLEVs were separated from soluble PLAP by centrifugation and NDOG2-nanobeads used to pull STB-EVs from total plasma EVs. Expression of miR518 (from the placental chromosome 19 microRNA cluster) was used to demonstrate assay sensitivity (Figure 4B). Perfusion STB-EVs were spiked into non-pregnant plasma as a positive control, achieving around 2000-fold greater miR518 expression than pregnant plasma. Nanobeads without antibody (saturated with free biotin) were added to pregnant plasma as a negative control; no miR518 expression was detected.

This technique was used to quantify the abundance of 5’-tRF-Glu-CTC (tRF-A) in peripheral plasma from 14 women with early-onset preeclampsia and 12 gestation-matched normotensive controls (Supplementary Table 5). 5’-tRF-Glu-CTC was upregulated (median 1.4-fold, p=0.017) in preeclampsia plasma STB-EVs (Figure 4C). The effect size was comparable to preeclampsia perfusion STB-EVs (median 1.4-fold upregulated).

**Figure 4.**
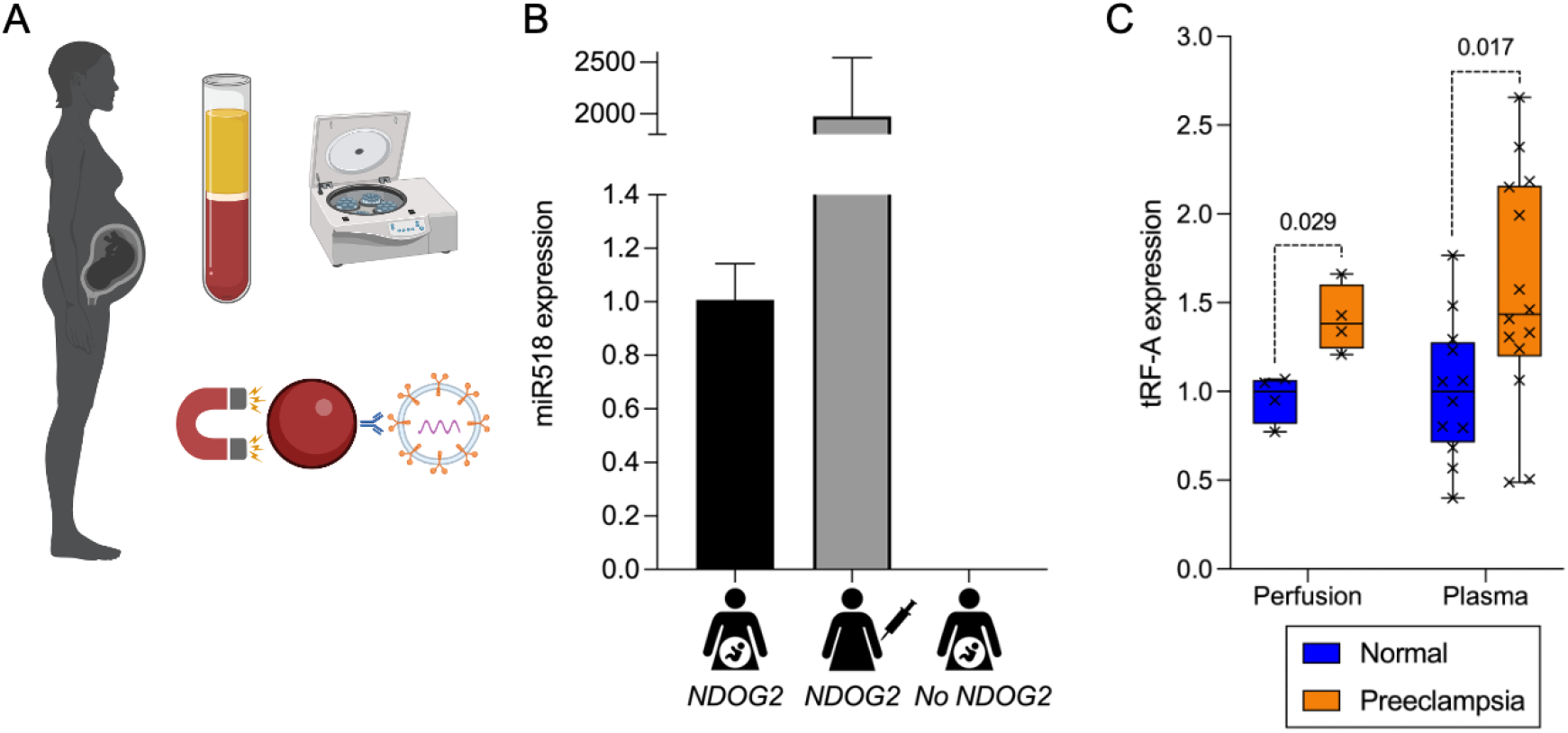
Isolation of STB-EVs from venous plasma A: Diagram summarising protocol for STB-EV isolation: plasma preparation, 10 000 *g* centrifugation to isolate MLEVs, immuno-isolation using magnetic beads coated with antibody to placental alkaline phosphatase (NDOG2). B: Relative expression of miR518 in EVs isolated from three samples using above technique (from left to right: pregnant plasma using beads coated with NDOG2; non-pregnant plasma spiked with perfusion STB-EVs using beads coated with NDOG2; pregnant plasma using beads without NDOG2 coating). Bars represent median, error bars represent interquartile range. C: Relative expression of 5’-tRF-Glu-CTC (tRF-A) in perfusion-derived STB-EVs (not gestation-matched, reproduced from Figure 2B for comparison) and gestation-matched peripheral venous plasma STB-EVs from women with early-onset preeclampsia (n=14) and normotensive pregnancies (n=12).

### EV-bound 5’-tRF-Glu-CTC promotes macrophage, but not monocyte activation

Inflammation is a key feature of preeclampsia^27^. Tissue-resident macrophages are known to be activated in preeclampsia^28^. Recent studies suggest STB-EVs may underlie this activation^29^. We treated macrophages and monocytes in culture with the most abundant 5’-tRF upregulated in preeclampsia (5’-tRF-Glu-CTC, labelled A for brevity; full sequence in Supplementary Table 2). RNA was packaged within otherwise undecorated lipid vesicles, in order to distinguish the effect of one tRF from the accompanying RNA, lipids and proteins in perfusion-derived STB-EVs. We used a scrambled version of the most abundant STB-EV 5’-tRF sequence as a negative control, following validation against untreated cells (Supplementary Figure 2). Macrophages were activated to a type 1 phenotype after 12 hours treatment with tRF-A, increasing expression of pro-inflammatory cytokines (Figure 5A). This pro-inflammatory 5’-tRF action was confined to macrophages, with no changes observed when the same experiment was repeated in undifferentiated monocytes (Figure 5B).

**Figure 5.**
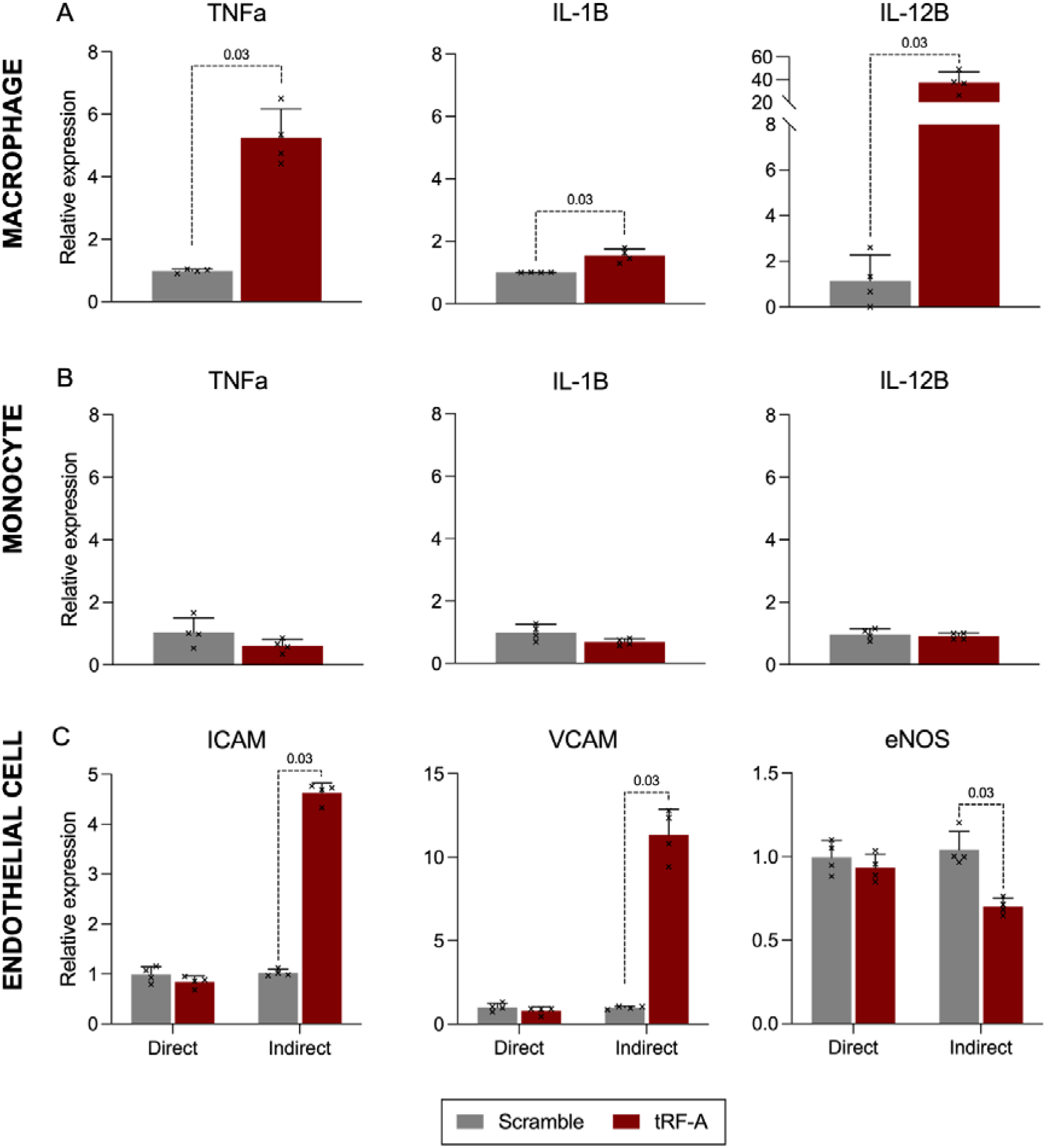
5’-tRF-Glu-CTC (tRF-A) actions in immune and endothelial cells. A: Cytokine expression by THP-1 derived macrophages after treatment with vesicle-encapsulated RNA for 12 hours. B: Cytokine expression by THP-1 monocytes (undifferentiated) after treatment with vesicle-encapsulated RNA for 12 hours. C: Expression of cell adhesion molecule and endothelial nitric oxide synthase mRNA in human umbilical vein endothelial cells treated with vesicle-encapsulated RNA (Direct), or the cell culture medium from macrophages pre-treated with vesicle-encapsulated RNA (Indirect) for 6 hours. Fragments labelled tRF-A and Scramble for brevity; full sequences in Supplementary Table 2 and Methods; n=4 per condition.

### EV-bound 5’-tRF-Glu-CTC indirectly activates endothelial cell adhesion molecules and reduces expression of endothelial nitric oxide synthase

Endothelial damage is considered a common factor in the wide-ranging maternal organ dysfunction of preeclampsia^1^. Vascular macrophages are known to modulate adjacent endothelial cell function^30^. We treated human umbilical vein endothelial cells with vesicle-bound tRF-A (direct) and with the supernatants from tRF-A-activated macrophages (indirect) alongside corresponding scramble controls. tRF-A activated endothelial cells (quantified through increased expression of adhesion molecules) indirectly through macrophage stimulation, but not directly (Figure 5C).

The primary presenting feature of preeclampsia is high blood pressure. Immediate blood pressure regulation involves constitutive release of nitric oxide from endothelial cells, which maintains a state of relative vasodilation. In preeclampsia, circulating nitrite levels are reduced, which may contribute to hypertension^31^. tRF-A reduced expression of endothelial nitric oxide synthase in endothelial cells, indirectly via macrophage activation (Figure 5C).

## Discussion

We are the first to report 5’-tRFs as the predominant species of small RNA in preeclampsia STB-EVs. These data are consistent with our prior finding in healthy pregnancy STB-EVs^11^. Enrichment of 5’-(but not 3’-) tRFs in EVs is also reported in immune cells and suggests a specific export process^13,17^. We demonstrate differential expression of over 900 STB-EV small RNA fragments in early-onset preeclampsia compared to normal pregnancy. Different preeclampsia phenotypes are unified by stress in the syncytiotrophoblast^3^; tRFs are produced by stress-dependent ribonucleases^12^. Thus, a change to STB-EV 5’-tRF expression fits with our existing understanding of preeclampsia. Differential expression in preeclampsia was identified in MLEVs but not SEVs, which is consistent with different EV biogenesis: SEVs are released constitutively via the endosomal pathway whereas MLEVs form by budding in response to stress^32^. The effect size between preeclampsia and normal is consistent with other studies of differential 5’-tRF expression in disease^33^. We corroborated discoveries in perfusion data by finding increased STB-EV 5’-tRF-Glu-CTC in preeclampsia plasma compared to normotensive controls.

One of the most studied tRNA ribonucleases is angiogenin, which is reported to generate 2-3 phosphate residues at the 3’ end of the 5’-tRF. Our sequencing approach detected 5’-tRF with hydroxyl groups, but not 2-3 cyclic phosphate residues, suggesting STB-EV 5’-tRFs were generated by ribonucleases other than angiogenin^34^.

5’-tRFs are known to influence cellular function through a variety of regulatory mechanisms at the level of the transcriptome, translatome and the epigenome^15^. We found 5’-tRF-Glu-CTC directly activated macrophages, but not monocytes or umbilical vein endothelial cells. We speculate this difference may be accounted for by phagocytosis of EVs by macrophages, trafficking 5’-tRFs to the endosomal compartment (usually free of nucleic acids) where they could encounter Toll-like receptor 7. This hypothesis is founded in published work demonstrating a lack of macrophage response to unencapsulated 5’-tRFs, or 5’-tRFs with Toll-like receptor 7 antagonists and warrants further investigation^17^. Macrophages are not typically in direct contact with blood; however in preeclampsia the endothelial barrier is significantly disrupted^1^. We suggest that circulating EV-bound 5’-tRFs would directly reach macrophages in the vessel walls in preeclampsia, where their functional effect could contribute to the well-described sterile inflammation of the disease^27^. Our findings of increased pro-inflammatory cytokine expression in response to tRF-Glu-CTC correlate with plasma cytokine concentrations in preeclampsia^35^.

STB-EVs are known to directly damage the endothelium, yet we found no direct effect of 5’-tRF-Glu-CTC on human umbilical vein endothelial cells^36^. This discrepancy may be attributed to the absence of other EV RNA and proteins which could be necessary to trigger endothelial damage. Our data show 5’-tRF-Glu-CTC macrophage activation indirectly activates the endothelium.

Prior studies of 5’-tRFs have used cell culture as a model system. Work in breast cancer reported intracellular 5’-tRF expression promoted metastasis^16^. Mycobacterium infection in human macrophages triggered 5’-tRF release in EVs, activating neighbouring cells^17^. Here we have investigated 5’-tRFs at a whole-organ level: the placenta is expelled with the fetus during parturition and can be studied intact *ex-vivo*. Sampling of the uterine vein during caesarean section has offered *in-vivo* evidence for a placental 5’-tRF load in maternal plasma. An immuno-assay has corroborated STB-EV 5’-tRF-Glu-CTC upregulation in preeclampsia plasma. Together with functional data showing 5’-tRF-Glu-CTC macrophage activation, we propose a possible endocrine signalling function for 5’-tRFs, contributing to preeclampsia pathogenesis.

Our study’s strength lies in the unique integration of cutting-edge techniques and distinctive samples. Previous studies investigating STB-EV small RNA have used lower fidelity models to source EVs (e.g. explants) and taken bioinformatic approaches which disregard 5’-tRF data, despite noting their presence^37,38^. We have corroborated perfusion-based RNA sequencing discoveries *in vivo* using qPCR in plasma. We overcame confounding in high-profile studies of total cell-free RNA in preeclampsia by focussing our attention exclusively on placental RNA^39,40^. The smaller size of our discovery cohort could be considered a weakness in a heterogeneous disease; we consider that by confining ourselves to a common step (syncytiotrophoblast stress) in early-onset disease, as well as confirming our data *in vivo* and *in vitro*, our findings are pertinent.

### Perspectives

Preeclampsia is a multifactorial condition, with diverse clinical features. Syncytiotrophoblast stress is common to all cases, but remains poorly understood. Here we present 5’-tRFs, a novel and highly abundant class of RNA differentially released by the preeclamptic syncytiotrophoblast. We demonstrate placentally-derived 5’-tRFs in the maternal circulation. The most abundant preeclampsia-upregulated STB-EV 5’-tRF was also increased in preeclampsia plasma. We find pro-inflammatory effects of this tRF on macrophages. Together these data suggest 5’-tRFs may play a role as transducers of an inflammatory signal from placenta to periphery in preeclampsia. Our findings offer a novel category of signalling molecule released by the placenta, warranting further investigation. We speculate that 5’-tRFs may dysregulate other cells in preeclampsia. Future studies will consider: other putative 5’-tRF targets in preeclampsia, such as liver sinusoids and pericytes; the actions of additional 5’-tRFs which are differentially expressed in preeclampsia; whether STB-EV 5’-tRFs play a role in other pregnancy-related diseases. Our ongoing work also focuses on the optimisation of techniques to isolate low-abundance placental EV 5’-tRF signal from complex biofluids such as plasma, with attention to their clinical relevance. 5’-tRFs may join other better-studied stress markers such as sFlt-1 and PlGF in explaining the pathogenesis of preeclampsia.

### Novelty and Relevance

#### 1. What is New?

5’-tRFs are the most abundant small RNA species within preeclampsia STB-EVs and are differentially expressed compared to normal pregnancy. A proportion of 5’-tRFs in the maternal circulation are placentally-derived. The most abundant upregulated 5’-tRF in preeclampsia STBEVs is 5’-tRF-Glu-CTC; this has been discovered by placental perfusion and corroborated in plasma. 5’-tRF-Glu-CTC has pro-inflammatory effects on macrophages.

#### 2. What is Relevant?

Preeclampsia is a placentally-derived hypertensive disorder, which can result in multi-organ failure. Recent studies report STB-EVs could promote preeclampsia through macrophage activation. Our findings suggest 5’-tRFs may underlie some pro-inflammatory actions of STB-EVs in preeclampsia.

#### 3. Clinical/Pathophysiological Implications?

STB-EV 5’-tRFs represent a feto-maternal signal with sufficient complexity to contribute to the varied clinical features of preeclampsia. 5’-tRFs may represent a biomarker or therapeutic target in this syndrome.

## Acknowledgements

We wish to thank Fenella Roseman, research midwife, for assistance with collecting clinical samples.

## Sources of Funding

WRC receives a clinical doctoral training fellowship from the Wellcome Trust, which funded this study.

## Disclosures

YMDL holds equity in DRA, Insighta Grail/Illumina and Take2. PJ holds equity in Illumina. PJ is a consultant to Take2. PJ is a Director of DRA and KingMed Future. YMDL, PJ, and LJ receives royalties from Illumina, LabCorp, Grail, DRA, Xcelom and Take2.

## Author contributions

Study was conceived and designed by WRC, CR and MV. Bioinformatic analyses were performed by PJ, LJ, JB, WRC and GDJ. Supervision was provided by PJ, YMDL, CR and MV. Experimental work was performed and manuscript written by WRC. All authors edited and approved the final manuscript.

## Abbreviations

DOTAP: N-[1-(2,3-Dioleoyloxy)propyl]-N,N,N-trimethylammonium methylsulfate EV Extracellular vesicle
MLEV: Medium-large extracellular vesicle PlGF Placental growth factor
SEV: Small extracellular vesicle
STB-EV: Syncytiotrophoblast extracellular vesicle sFlt-1 Soluble fms-like tyrosine kinase-1
Trf: tRNA fragment

## Notes

### Summary of Updates

Introduced plasma data quantifying 5'-tRF-Glu-CTC in plasma STB-EVs in new figure (Figure 4) and in abstract, methods, results, discussion, perspectives, novelty and relevance (in response to reviewers' comments). Other small changes in light of novel data. Reformatted.

