## Supplementary Figures and Tables for "Differential Expression of 5’-tRNA Fragments in Circulating Preeclampsia Syncytiotrophoblast Vesicles Drives Macrophage Inflammation"

Supplemental Material

*Supplementary Figure 1*

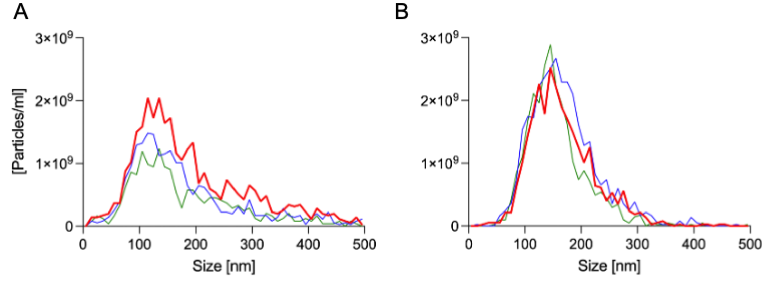

Nanoparticle Tracking Analysis size profiles for three separate samples for:

A) STB-MLEVs obtained by placental perfusion.

B) EVs obtained from eluate of membrane affinity columns loaded with maternal plasma.
*Supplementary Figure 2*

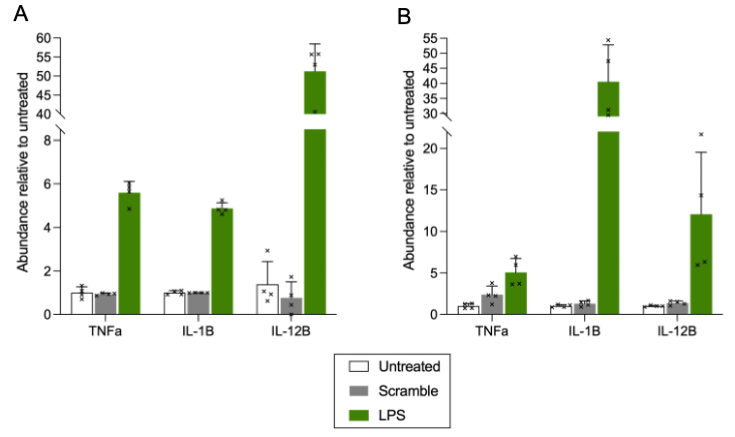

Scramble RNA control has no notable effect on macrophage cytokine expression when compared to untreated cells. Lipopolysaccharide (LPS) used as a positive control.
*Supplementary Table 1*

|  | **Normal (n=6)** | **Preeclampsia (n=8)** | **P value** |
| --- | --- | --- | --- |
| Median maternal age (IQR) – years | 34.5 (30.5-38.75) | 32 (30.3-37.5) | 0.59 |
| Median pre-pregnancy BMI (IQR) | 27 (20.8-33.8) | 28.3 (23.8-38) | 0.43 |
| First pregnancy – no. (%) | 0 (0) | 4 (50) | 0.08 |
| Diabetes – no. (%) | 0 (0) | 0 (0) | >0.99 |
| Essential hypertension – no. (%) | 0 (0) | 2 (25) | 0.47 |
| Current smoker – no. (%) | 0 (0) | 0 (0) | >0.99 |
| Male fetus – no. (%) | 4 (67) | 4 (50) | 0.63 |
| Median birthweight (IQR) – g | 3775 (3419-4090) | 1398 (1110-2270) | 0.001 |
| Median gestational age at delivery (IQR) – weeks | 39.2 (38.8-40) | 31.4 (30-35.3) | 0.0007 |
| Median highest recorded SBP (IQR) – mmHg | 132 (113-135) | 164 (140-178) | 0.003 |
| Median highest recorded DBP (IQR) – mmHg | 74 (64-75) | 108 (94-112) | 0.003 |

Characteristics of women who donated their placentas for dual-lobe perfusion, described using median/interquartile range (IQR) and compared using unpaired, two-sided Mann-Whitney tests; or described using number (no.) and percentage (%) and compared using two-sided Fisher’s exact tests.
*Supplementary Table 2*

| **ID** | **Sequence** | **Length** | **Mapping no indel** | **Mapping ≤2 indel** | **Mean PE count** | **SD PE count** | **Mean N count** | **SD N count** | **Log2Fold Change** | **Adjusted p value** | **Max monocyte** | **Max erythrocyte** | **Max platelet** |
| --- | --- | --- | --- | --- | --- | --- | --- | --- | --- | --- | --- | --- | --- |
| A | TCCCTGGTGGT  CTAGTGGTTAG  GATTCGGCGCT | 33 | tRF-Glu-CTC | tRF-Glu-CTC | 52051 | 18436 | 18405 | 5423 | -1.500 | 0.0054 | 204.5 | 43.3 | 10 |
| B | GCATTTGTGGT  TCAGTGGTAGA  ATTCTCGCCTG | 33 | tRF-Gly-GCC | tRF-Gly-GCC | 1159 | 296 | 534 | 193 | -1.117 | 0.0225 | 6.2 | 0.8 | 2 |
| C | CCCCTGTGGTC  TAGTGGTTAGG  ATTCGGCGCC | 32 | Unmatch | tRF-Glu-CTC | 965 | 124 | 530 | 95 | -0.865 | 0.0058 | 0.9 | 0 | 0 |

Median normalised expression of three target 5’-tRFs (labelled A, B and C for brevity) in placental EVs and maximum normalised expression of the same 5’-tRFs in blood-cell datasets. All count values represent reads per million (RPM).
*Supplementary Table 3*

| **Sample** | **SRA ID** |
| --- | --- |
| Monocyte_1 | SRR6453428 |
| Monocyte_2 | SRR6453427 |
| Monocyte_3 | SRR6453426 |
| Monocyte_4 | SRR6453386 |
| Monocyte_5 | SRR6453387 |
| Monocyte_6 | SRR6453388 |
| Erythrocyte_1 | SRR1664893 |
| Erythrocyte_2 | SRR1664894 |
| Erythrocyte_3 | SRR1664895 |
| Erythrocyte_4 | SRR1664896 |
| Erythrocyte_5 | SRR1664897 |
| Platelet_1 | SRR10282877, SRR10282878 |
| Platelet_2 | SRR10282879, SRR10282880 |

Sequence read archive (SRA) identifiers for blood cell datasets

*Supplementary Table 4*

| **Assay type** | **Manufacturer** | **Target** | **Identifier** |
| --- | --- | --- | --- |
| TaqMan™ gene expression assay | Applied Biosystems, USA | *TBP* | Hs00188166_m1 |
|  |  | *GAPDH* | Hs02758991_g1 |
|  |  | *ICAM1* | Hs00164932_m1 |
|  |  | *VCAM1* | Hs01003372_m1 |
|  |  | *NOS3* | Hs01574665_m1 |
|  |  | *TNF* | Hs00174128_m1 |
|  |  | *IL1B* | Hs01555410_m1 |
|  |  | *IL12B* | Hs01011518_m1 |
| TaqMan™ MicroRNA Assay | Applied Biosystems, USA | cel-miR-39 | 000200 |
|  |  | miR518 | 001156 |

Quantitative polymerase chain reaction assays used for RNA detection

*Supplementary Table 5*

|  | **Normal (n=12)** | **Preeclampsia (n=14)** | **P value** |
| --- | --- | --- | --- |
| Maternal age in years – median (IQR) | 30 (28-33) | 30 (26-35) | 0.69 |
| Pre-pregnancy BMI – median (IQR) | 24 (21-27) | 23 (22-25) | 0.80 |
| First pregnancy – no. (%) | 6 (50) | 11 (79) | 0.22 |
| Diabetes – no. (%) | 0 (0) | 0 (0) | >0.99 |
| Essential hypertension – no. (%) | 0 (0) | 0 (0) | >0.99 |
| Current smoker – no. (%) | 2 (17) | 0 (0) | 0.22 |
| Male fetus – no. (%) | 7 (58) | 5 (36) | 0.43 |
| Gestational age at sample in weeks – median (IQR) | 31.9 (30-33.4) | 31.3 (28.8-32.7) | 0.64 |
| Birthweight in grams – median (IQR) | 3180 (2943-3291) | 1623 (1103-1766) | <0.0001 |
| Gestational age at delivery in weeks – median (IQR) | 39.1 (37.5-39.8) | 32.1 (29.7-34.2) | <0.0001 |
| Highest recorded SBP in mmHg – median (IQR) | 131 (120-134) | 180 (160-180) | <0.0001 |
| Highest recorded DBP in mmHg – median (IQR) | 74 (70-80) | 110 (98-115) | <0.0001 |
| Days from diagnosis to delivery – median (IQR) | N/A | 9 (6-18) | - |

Characteristics of women who donated blood samples, described using median and interquartile range (IQR) and compared using unpaired, two-sided Mann-Whitney tests; or described using number (no.) and percentage (%) and compared using two-sided Fisher’s exact tests.
